# Repurposing the biased visibility in HiC datasets to mark dynamically regulated condensed and decondensed chromatin states genome-wide

**DOI:** 10.1101/498006

**Authors:** Keerthivasan Raanin Chandradoss, Prashanth Kumar Guthikonda, Srinivas Kethavath, Monika Dass, Harpreet Singh, Rakhee Nayak, Sreenivasulu Kurukuti, Kuljeet Singh Sandhu

**Affiliations:** Department of Biological Sciences, Indian Institute of Science Education and Research (IISER) – Mohali Knowledge City, Sector 81, SAS Nagar 140306, India; Department of Animal Biology, School of Life Sciences, University of Hyderabad (UoH), Prof. CN Rao Road, P O, Central University Gachibowli, Hyderabad, Telangana 500046, India

**Keywords:** HiC, 3D genome, chromatin condensation, Lamina associated domains, CTCF

## Abstract

Proximity ligation based techniques, like HiC, involve restriction digestion followed by ligation of formaldehyde cross-linked chromatin. Through analysis of lamina-associated domains (LADs), inactive X-chromosome in mammals and polytene bands in fly, we first established that the DNA in condensed chromatin had lesser accessibility to restriction endonucleases used in HiC as compared to that in decondensed chromatin. The observed bias was independent of known systematic biases, was not appropriately corrected by existing computational methods, and needed an additional optimization step. We then repurposed this bias to identify novel condensed domains outside LADs, which were bordered by insulators and were dynamically associated with the developmentally regulated epigenetic and transcriptional states. Our observations suggested that the corrected one-dimensional read counts of existing HiC datasets can be reliably repurposed to study the gene-regulatory dynamics associated with chromatin condensation and decondensation.

## Introduction

The three-dimensional genome organization is tightly linked with the regulation of essential genomic functions like transcription, replication and genome integrity(Bonev and Cavalli, 2016; Dekker and Mirny, 2016; Garcia-Nieto et al., 2017; Gonzalez-Sandoval et al., 2015; Therizols et al., 2014). While the significance of genome organization has been realized several decades ago, the comprehensive evidence emerged somewhat recently through the advent of proximity ligation based techniques like Chromosome Conformation Capture (3C), Circular-3C (4C,) 3C-Carbon-Copy (5C) and HiC(Dekker et al., 2002; Dostie et al., 2006; Lieberman-Aiden et al., 2009; Simonis et al., 2006; Zhao et al., 2006). It is recognized that the eukaryotic genome is hierarchically organized into self-interacting topologically associated domains (TADs), which can have distinct chromatin states that were insulated from neighbourhood through boundaries marked with CCCTC-binding factor (CTCF), Cohesins, ZNF143 and Top2b factors (Dixon et al., 2016; Dixon et al., 2012; Heidari et al., 2014; Uuskula-Reimand et al., 2016). The TADs are ancient genomic features and are depleted for evolutionary breakpoints within (Harmston et al., 2017; Krefting et al., 2018). Fudenberg et al proposed that that chromatin extrudes through the ring formed by the Cohesins until the chromatin encounters the CTCF insulator, a model known as ‘loop extrusion’ model (Fudenberg et al., 2016). CTCF binding is transiently lost during pro-metaphase, which coincides with the loss of TAD structures during M-phase (Agarwal et al., 2017; Nagano et al., 2017) (Oomen et al 2018, biorxiv). Systematic depletion of CTCF and Cohesins also leads to de-insulation and partial disruption of TADs (Nora et al., 2017; Schwarzer et al., 2017). An array of studies has shown that TADs function as basic units of 3D genome that dynamically associate with the epigenetic states of genes, including replication timing, during development and differentiation (Bonev et al., 2017; Boya et al., 2017; Flyamer et al., 2017; Fraser et al., 2015; Kaaij et al., 2018; Ke et al., 2017; Le Dily et al., 2014; Neems et al., 2016; Pope et al., 2014). How these dynamical epigenetic states of TADs are regulated is not entirely clear. One of the ways, this can be achieved is through chromatin condensation and decondensation, implying inactive and active states of TADs respectively (Chambeyron and Bickmore, 2004; Ciabrelli and Cavalli, 2015; Rafique et al., 2015; Therizols et al., 2014; Williamson et al., 2014) (Benabdallah et al 2018, Biorxiv). While it is established that the gene-poor and transcriptionally inactive domains locate towards nuclear periphery, and remain stably condensed during differentiation with local gene-specific alterations(Peric-Hupkes et al., 2010), the dynamics of chromatin condensation and decondensation in the other regions of the genome largely remains under-explored. Condensation and decondensation of chromatin is generally studied through microscopic methods. In this study, we demonstrated that the condensed and decondensed states of chromatin domains could be directly identified from one dimensional HiC read counts.

Yaffe and Tanay have shown that HiC datasets have systematic bias due to differential ligation efficiency of restriction fragments of different lengths, the differential amplification of fragments with GC rich ends and differential mappability of sequences(Yaffe and Tanay, 2011). Several methods have since been developed to normalize the aforementioned biases. These methods can be broadly categorized into two classes, the ones that define the aforementioned biases explicitly in the algorithm and the ones which do not define the source of bias and instead adopt an implicit approach based on fractal folding of the chromatin and the equal visibility of all genomic loci(Cournac et al., 2012; Hu et al., 2012; Imakaev et al., 2012; Rao et al., 2014; Yaffe and Tanay, 2011). We showed that the differential visibility of genomic loci to the restriction endonucleases used in HiC protocols caused potential bias in HiC data. HiC reads were significantly depleted for the interactions impinging from condensed heterochromatin domains, and this bias was not appropriately corrected by existing computational methods. By repurposing the observed bias, we first demonstrated that the bias in one-dimensional read counts of HiC datasets reliably marked the known condensed and decondensed domains in the genome and then highlighted the developmentally regulated dynamics of condensed and decondensed states of chromatin.

## Results

### Biased visibility in HiC data marks condensed and decondensed chromatin domains

We first showed that the *in-situ* restriction digestion of chromatin was not uniform in the genome. Towards this, we obtained the sequencing data for *in-situ* restriction digested chromatin and *in-solution* restriction digested naked DNA of mouse embryonic stem cells (mESC) (Chen et al., 2014). We calculated the read counts for 10 kb bins of the mouse genome and normalized by the total reads. We further corrected the read counts for restriction site density (RE-density) of the bins and the GC content of the bins using loess regression, in that order (Methods, Figure S1a-b). The scatter-plot of restriction digested naked DNA and *in-situ* digested chromatin showed skew towards naked DNA axis marking the inefficient digestion of certain genomic regions in chromatin but not in naked DNA (Figure 1a). This suggested that chromatin structure had influenced its own digestibility. The likely explanation was that the decondensed chromatin was readily digested while heterochromatin domains had limited accessibility to restriction endonuclease. To assess this hypothesis, we obtained the Lamina Associated Domains (LADs), which are known heterochromatin domains attached to the nuclear periphery in condensed form(Ciabrelli and Cavalli, 2015; van Steensel and Belmont, 2017). We calculated the raw and corrected read counts in the constitutive LADs (cLADs) and constitutive inter-LADs (ciLADs) in mESC. As shown in the figure 1b-c, cLADs exhibited significantly less raw read counts as compared to ciLADs in *in-situ* digested chromatin as well as in *in-solution* digested naked DNA, suggesting that the reads from digested naked DNA had bias likely due to varying densities of restriction sites and the relative GC content of cLADs and ciLADs (Figure 1b-c, p<2.2e-16). The read counts corrected for RE-density and GC content, however, exhibited bias only in the *in-situ* digested chromatin and not in the naked DNA, highlighting that the cLADs were relatively inaccessible to restriction endonuclease likely due to condensed nature of the chromatin (Figure 1b-c, p<2.2e-16). We further identified the chromatin domains significantly enriched (decondensed) or depleted (condensed) in corrected read counts (Methods, Figure S1c). Overall, 77% of length covered by condensed domains were within cLADs and 23% mapped to ciLAD regions, marking the condensed domains which were not associated with the nuclear lamina (Figure S1d).

**Figure 1.**
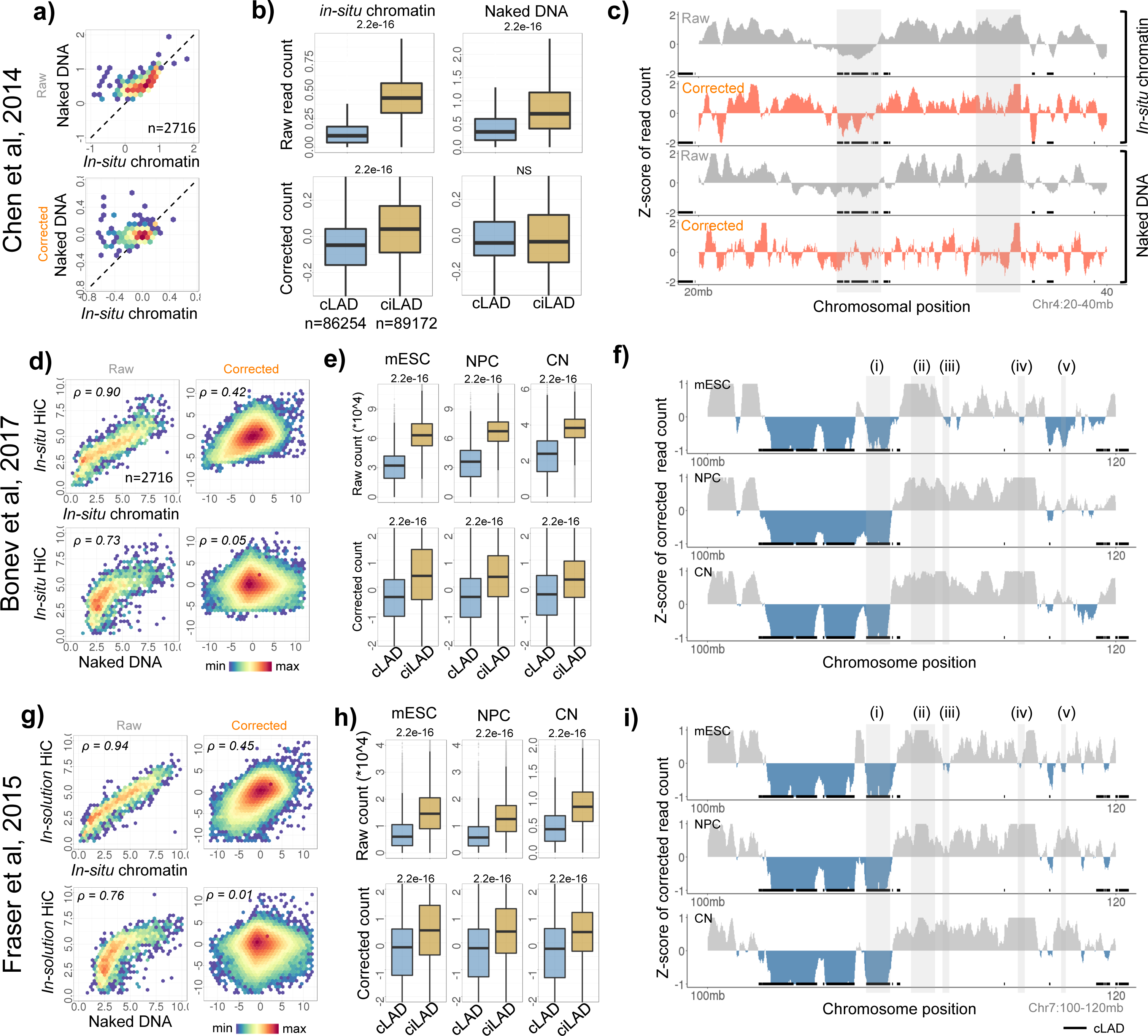
Biased visibility of chromatin domains in *in-situ* HiC datasets. (a) Scatter plots of raw and corrected read counts (per Mb) in *in-situ* digested chromatin vs. *in-solution* digested naked DNA. (b) Distribution of raw and corrected read counts of *in-situ* digested chromatin and *in-solution* digested naked DNA in cLADs and ciLADs. P-values were calculated using two-tailed Mann-Whiney U tests. (c) Illustrative example of raw and corrected read counts of *in-situ* digested chromatin and *in-solution* digested naked DNA along chr4: 20-40Mb region. (d) Top: scatter plots of raw and corrected read counts in *in-situ* HiC and *in-situ* digested chromatin. Bottom: scatter plots of raw and corrected read counts in *in-situ* HiC and *in-solution* digested naked DNA. ‘*ρ*’ represents the Pearson’s correlation coefficient. (e) Distribution of raw and corrected read counts of *in-situ* HiC datasets in cLADs and ciLADs. P-values were calculated using two-tailed Mann-Whiney U test. (f) Illustrative examples of corrected read counts of *in-situ* HiC datasets along chr7: 100-120 Mb. Regions (i) and (ii) mark constitutively condensed and decondensed regions respectively. Regions (iii)-(v) mark cell-type specific condensed and decondensed states. (ii) (g-i) Same as d-f, but for *in-solution* HiC data obtained from Fraser et al.

We then expanded our analyses to 40 HiC datasets (21 *in-situ* HiC, 11 *in-solution* HiC and 8 scHiC) and obtained the processed reads (Figure 1d-f, Figure S2, & Table S1). One-dimensional read counts were corrected for the density of restriction sites and GC content as earlier. Through analysis of mESC data, we observed that the reads from *in-situ* HiC had a significant correlation with the reads from *in-situ* digested chromatin, but exhibited skewed scaling towards digested naked DNA (Figure 1d). This suggested that the *in-situ* HiC reads exhibited bias similar to the one in *in-situ* digested chromatin. As shown in the figure 1e-f and figure S2, the corrected read counts exhibited enrichment in ciLADs and depletion in cLADs (p<2.2e-16). Again, 70% of length covered by the condensed domains were within cLADs and 30% was within ciLAD regions, marking the condensed domains other than LADs. (Figure S1d). Our observations with cLADs and ciLADs were consistent with different *in-situ* HiC datasets, including single-cell HiC, generated using distinct restriction enzymes (Figure 1d-i and S2, p<2.2e-16). We illustrated the examples of condensed domains that mapped to cLADs, to ciLADs and the ones that exhibited cell-type specificity in the figure 1f and S2.

We also showed that the observed association with the cLAD and ciLADs was not due to HiC-specific processing of the sequencing data. We observed the bias in reads simply processed through bowtie too (Figure S3a, p<2.2e-16, Methods). Further, the biased visibility was not the property of *in-situ* HiC only, but was also observed in *in-solution* HiC too (Figure 1g-I, S3b, p<2.2e-16). As shown in the figure S3c, the corrected reads from *in-situ* HiC exhibited good correlation with those from *in-solution* HiC in the same cell-type (mouse fetal liver) from the same study(Nagano et al., 2015). This suggested that the visibility bias was not affected by the method of ligation and that the source of bias was the difference in accessibility to the restriction endonucleases, not the difference in ligation.

HiCNorm, an explicit method of HiC correction, failed to remove the bias in the read counts, supporting that the observed bias is independent of known systematic biases of HiC data (Figure 2a & S4, p<2.2e-16). Iterative correction, an implicit method, normalized the read counts attributing to its intrinsic nature of polishing the HiC matrices for equal visibility of all loci without defining the bias at first place (Figure 2a & S4). Data obtained from Genome architecture mapping (GAM)(Beagrie et al., 2017), which directly obtains the co-localized DNA segments through large number of thin nuclear sections and does not involve any restriction and ligation steps, did not exhibit any bias in the read counts. By comparing GAM and ICE-corrected HiC data, we further observed that ICE merely lifted the background and the obscure signals in the contact matrices. In the process of lifting the obscure signals in the poorly digested condensed regions, ICE inadvertently lifted the long-range background interactions among condensed domains as shown in the figure 2b and S4. To address this, we proposed that the ICE-corrected HiC datasets needed a further distance dependent optimization of interaction frequencies. We termed this additional step as Distance Sorted Contact Optimization (DiSCO) and implemented it on raw, HiCNorm-corrected and ICE-corrected HiC matrices. As shown in in the figure 2b, the method corrected the distance dependent bias in interaction frequencies of condensed and decondensed domains. Though DiSCO corrected only the distance dependent bias when implemented alone on the raw data, it was able to balance the contact matrices for most of the biases when combined with the ICE. In particular, the long-range interactions of condensed domains, which were inadvertently lifted by ICE, were corrected by DiSCO, and the short-range interactions remained largely unaltered (Figure 2b-c & S4). Inclusion of DiSCO did not reintroduce the bias in the ICE-corrected 1D read counts, suggesting the overall suitability of the approach (Figure S4b). The comparison with the GAM matrices also showed sub-TAD structures and other types of interactions in the condensed domains (Figure 2c & S4c-e), which were clearly not captured by raw or any of the corrected HiC matrices, suggesting the inherent differences in the experimental techniques in deciphering the organization of condensed chromatin that are needed to be explored thoroughly in near future.

**Figure 2.**
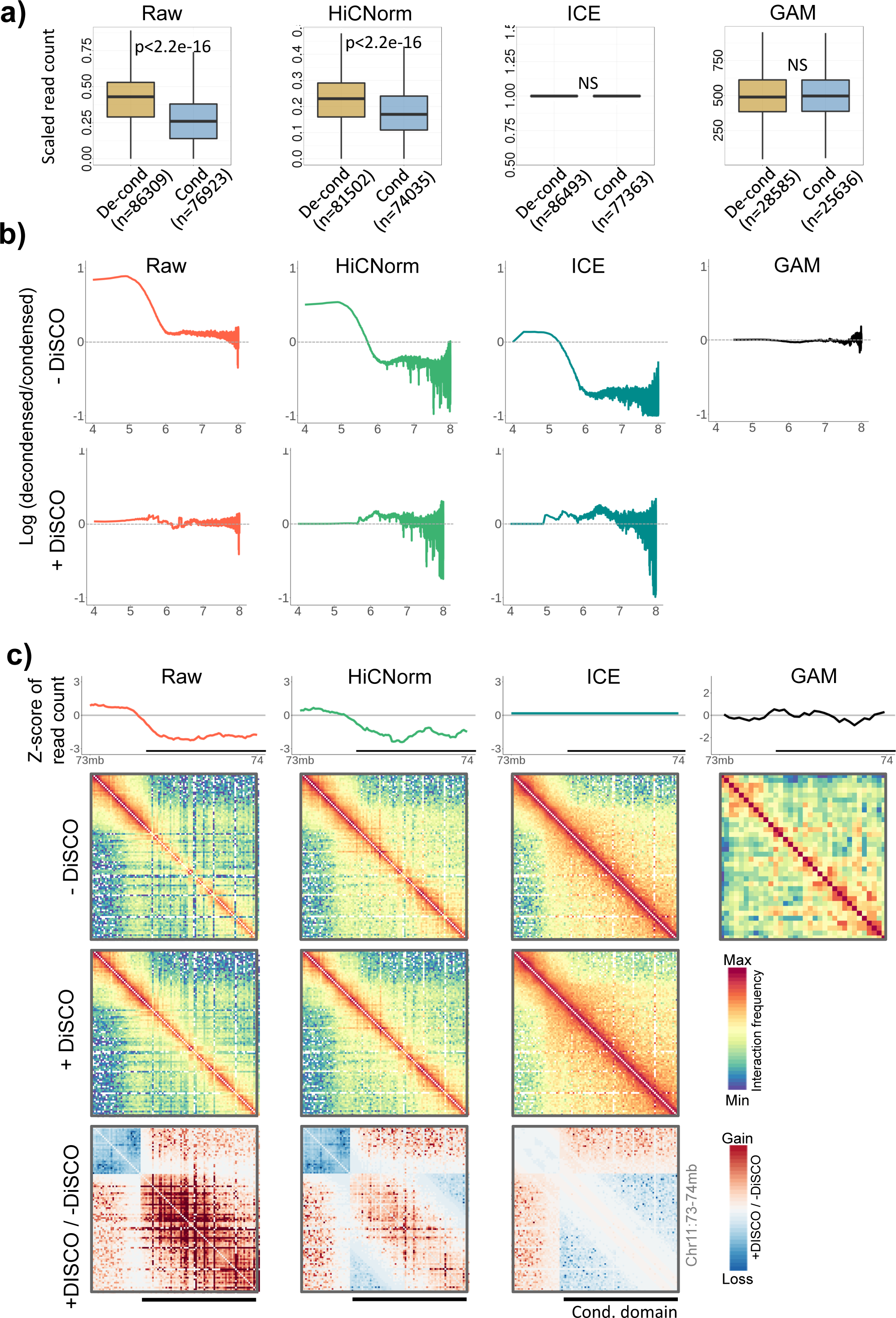
Bias in explicitly and implicitly normalized HiC, and GAM datasets. (a) Distribution of 1D read-counts of decondensed and condensed domains in raw, HiCNorm-corrected, ICE-corrected HiC and GAM datasets of mESCs. Values were scaled from 0 to 1. P-values were calculated using two-tailed Mann-Whiney U tests. (b) Upper panel: ratio of interaction frequencies of decondensed-to-decondensed and condensed-to-condensed interactions as a function of genomic distance in raw, HiCNorm-corrected, ICE-corrected and GAM datasets. Lower panel: plots after DiSCO correction. (c) Illustrative examples of raw, HiCNorm-corrected, and ICE-corrected data before and after DiSCO correction. Ratio matrices in the bottom panel show gain and loss of signals after DiSCO correction. GAM data is shown on extreme right for comparison. Additional examples are given in the Figure S4

To further scrutinize the differential digestion of condensed and decondensed domains, we obtained the HiC data for the polytene chromosome of Drosophila, which is a typical example of spatially condensed (polytene bands) and decondensed (inter-bands) domains (Eagen et al., 2015). The HiC reads were mapped and corrected as earlier. The analysis suggested that the polytene bands had lesser enrichment of corrected reads as compared to inter-band regions on both the polytene chromosome and the normal diploid chromosome (Figure 3a, p<2.2e-16). We illustrated our observations through examples in the figure 3b. On similar lines, we analysed the DNase-HiC data for active and inactive X-chromosomes in brain and patski cells (Deng et al., 2015). As shown through the scatter plots in figure 3c and examples in figure 3d, the X-chromosome had regions that were more visible in active X-chromosome and less visible in inactive X-chromosome. This suggested that the bias due to differential chromatin accessibility existed in both restriction endonuclease digested and DNase digested HiC datasets.

**Figure 3.**
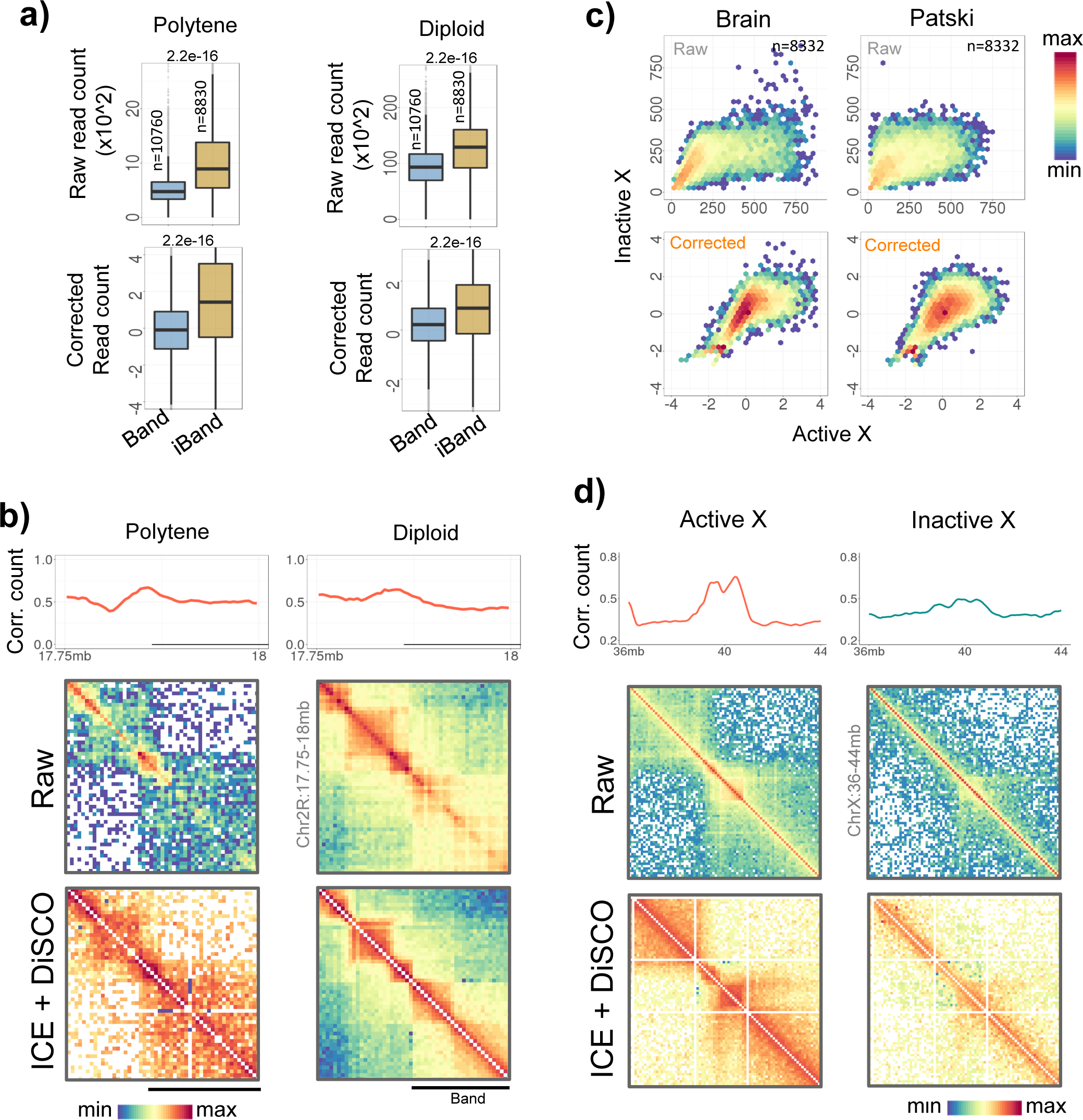
Low visibility of polytene bands and inactive X-chromosome. (a) Distribution of raw and corrected read counts in band and inter-band regions of polytene chromosome and the corresponding regions in diploid chromosome. P-values were calculated using two-tailed Mann-Whitney U tests. (b) Illustrative examples of read counts and contact maps in band and inter-band regions of polytene and diploid chromosome. Band regions are marked as horizontal line below the line plots. (c) Scatter plots of raw and corrected DNase-HiC read counts of active vs. inactive x-chromosomes in Brain and Patski cells. (d) Illustrative examples of corrected read counts and contact maps of chrX: 36-44 Mb region in active and inactive X-chromosome.

These observations highlighted that: 1) the observed bias in corrected 1D HiC read counts was independent of known systematic biases of HiC; 2) the bias captured the condensed and decondensed states of chromatin domains reliably, and 3) the existing computational approaches of HiC normalization needed further optimization for the condensed and decondensed domains.

### Dynamics of condensed and decondensed domains

To assess if the condensed and decondensed domains identified from restriction digestion bias in the ciLAD regions had functional significance, we analysed their dynamics during mouse embryonic stem cell (mESC) differentiation to neuronal progenitor cells (NPC) to cortical neurons (CN). As shown in the figure S5a, the differentiation from mESC to NPC exhibited greater overall change in corrected read counts as compared to NPC to CN differentiation. We, therefore, focussed on mESC to NPC differentiation to assess the developmental regulation of chromatin condensation and decondensation. We first mapped the histone modification and CTCF binding data around boundaries of domains by scaling all decondensed domains upstream and all condensed domains downstream to the domain boundaries (Figure 4a & S5b). We observed enrichment of active and inactive histone marks in decondensed and condensed domains respectively with transitions around boundaries that were marked with CTCF, RAD21, YY1, TOP2b, MIR and simple repeat elements (Figure 4a-b & S5c, p=4.5e-05 to 2.2e-16). Total 27.7% of condensed ciLAD domains in mESC were decondensed in NPC and 13.5% decondensed domains in mESC were condensed in NPC, suggesting significant cell-type specificity of domains identified through biased visibility in HiC data (Figure S5d). Genes exhibiting condensation during differentiation switched to repressed state and the ones showing decondensation switched to active state (Figure 4c, p=6.6e-13 & 2.2e-16). Through scatter plots of active and inactive chromatin types between mESC and NPC cells, we observed that the condensation of open chromatin domains during differentiation associated with the coherent change of active to inactive chromatin states (Figure 4d). Similarly, the domains that exhibited decondensation during differentiation switched to active states from inactive states (Figure 4d). Enrichment of neuronal development related terms among genes exhibiting decondensation, and the metabolism related terms among genes exhibiting condensation during ESC-to-NPC differentiation coherently supported the underlying change in their epigenetic and transcriptional states (Figure S6a). Figure 4e and S5e-g presented a few examples of chromatin domains that were constitutive (left panel) or cell-type specific (right panel) in mESC and NPC. These observations not only highlighted the developmental regulation of chromatin domains identified in the study, but also argued strongly against the dismissal of restriction digestion bias merely as an artefact.

**Figure 4.**
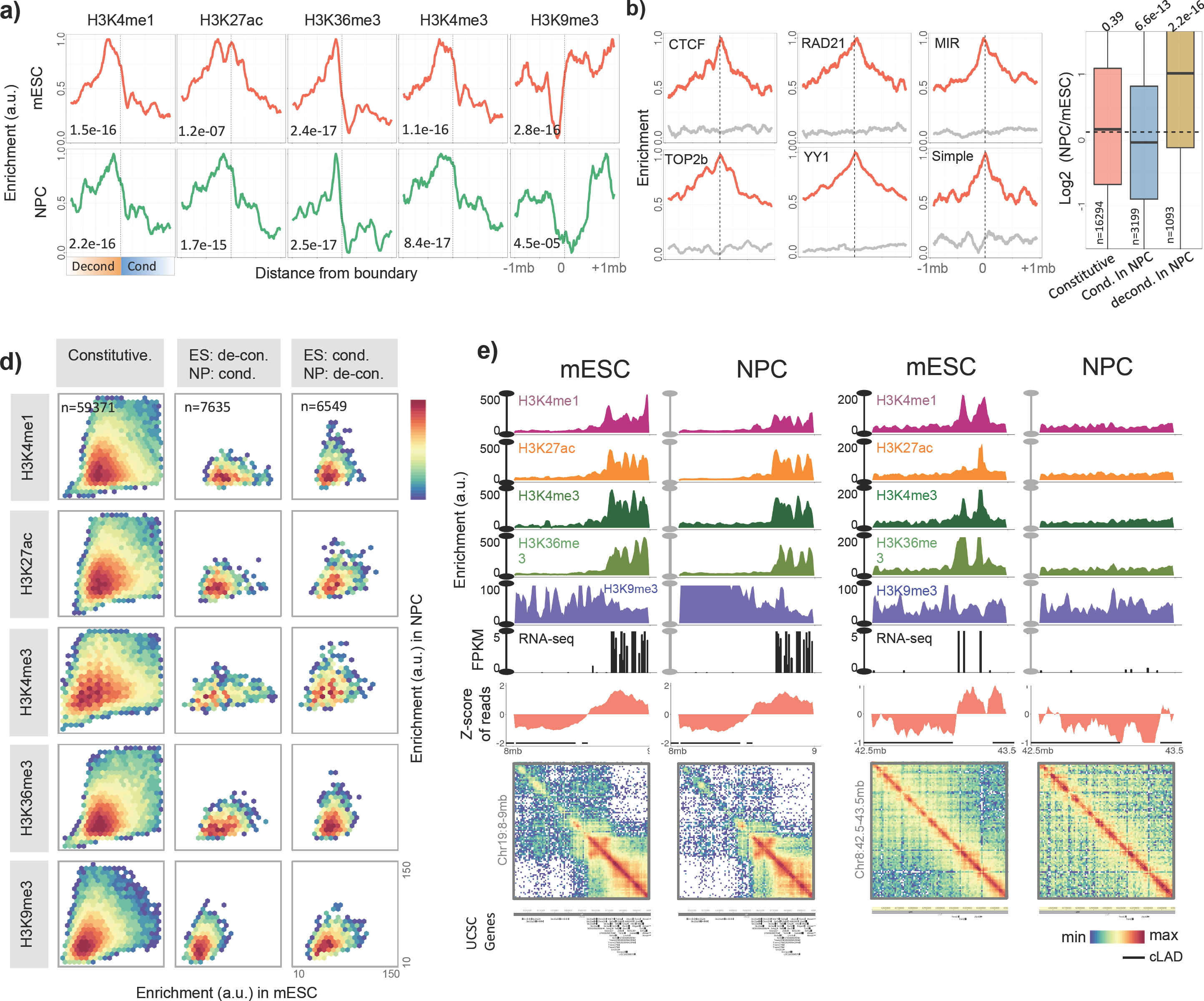
Developmental dynamics of chromatin condensation and decondensation. (a) Aggregation plots of histone modifications +/- 1Mb around the boundary of decondensed and condensed domains in mESC and NPC. P-values were calculated using two-tailed Mann Whitney U tests by comparing mean enrichment values in the bins of condensed and decondensed domains. (b) Enrichment of CTCF, RAD21, YY1, TOP2b binding, MIR and simple repeats +/- 1Mb around domain boundaries (red) and around domain centres (grey). (c) Boxplots representing change in gene expression in the chromatin domains that were constitutively present in mESC and NPC, the ones that switched to condensed state in NPC from decondensed state in mESC and *vice-versa*. P-values were calculated using two-tailed Mann-Whitney U tests of RPKM values in mESC and NPC. (d) Scatter plots of histone modifications in domains that remained unchanged in mESC and NPC, and the ones that switched from decondensed to condensed or *vice-versa* in mESC and NPC. (e) Examples of decondensed and condensed domains that remained consistent in mESC and NPC (left), and a decondensed region in mESC that switched to condensed state in NPC.

Chromatin condensation and decondensation can be induced by knocking out certain factors like Lamins. We, therefore, tested if such experimentally induced decondensation of LADs can be captured through analysis of HiC reads of Lamin knock out cells. We obtained the HiC data for WT and Lamin (*Lmb1, Lmb2, Lmna*) KO mouse embryonic stem cells from Zheng et al (Zheng et al., 2018). As shown in the figure 5a, the cLADs in Lamin KO cells exhibited relatively more reads as compared to that in WT (p<2.2e-16). We illustrated this observation through examples in figure 5b-c. Our observations highlighted that the HiC read-counts alone captured the decondensation of LAD domains after lamin deletion.

**Figure 5.**
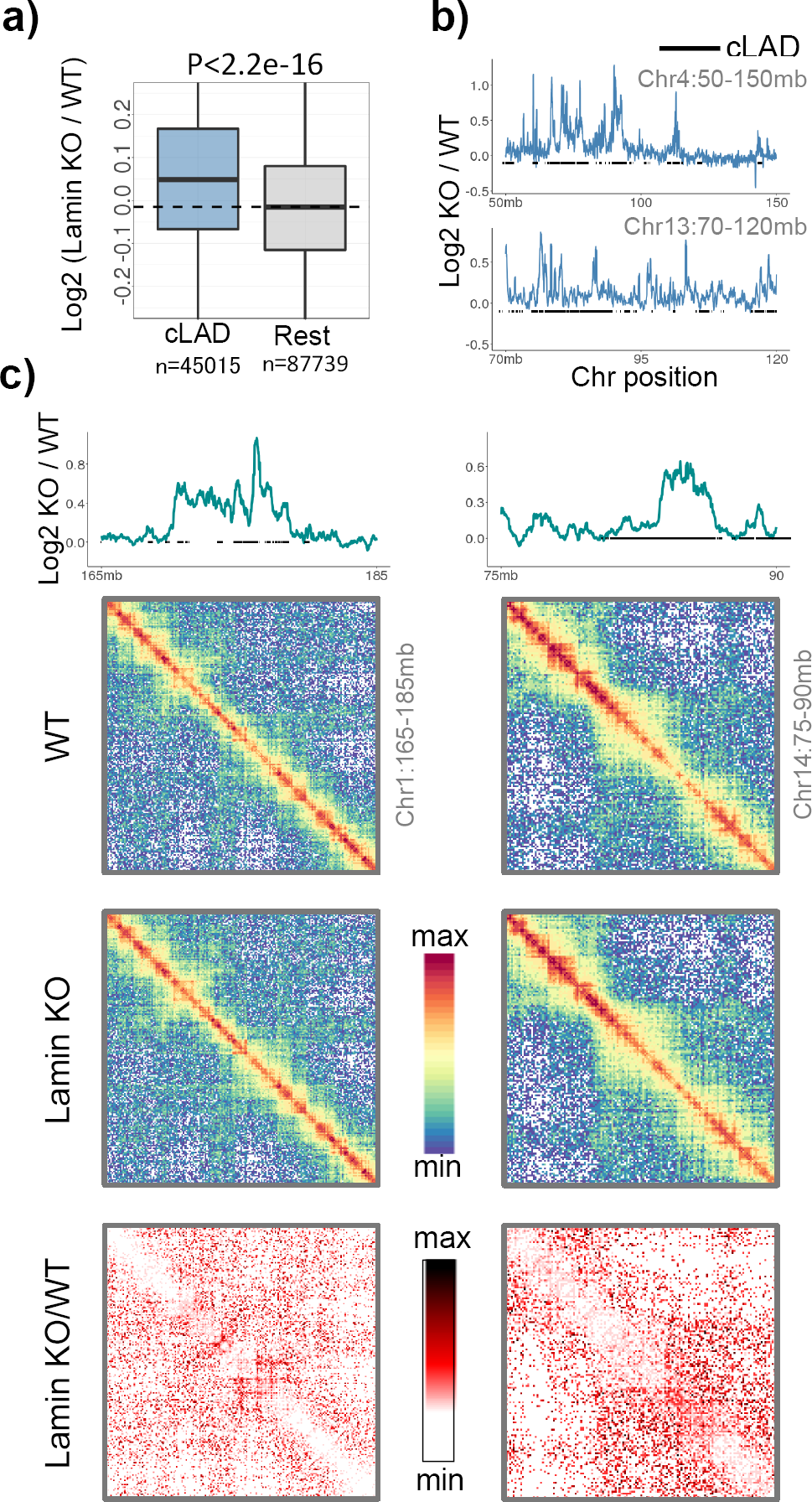
Capturing chromatin decondensation in Lamin KO cells. **(a)** Boxplots representing change in HiC read counts in the LAD domains after Lamin knock out in mESC cells. P-value was calculated using two-tailed Mann-Whitney U test. (b-c) Examples representing change in HiC read counts and contact matrices in WT and Lamin KO cells. The bottom panel of matrices represent the Lamin KO to WT fold-change in interaction frequencies.

## Discussion

To study the dynamics of gene regulatory activity in response to environmental and developmental clues, mapping chromatin accessibility remained an important task over decades. While DNase-I had been a preferred choice to digest the chromatin due to relatively lesser sequence specificity of the enzyme, only few studies had attempted restriction endonucleases to identify accessible regions in the chromatin (Chen et al., 2014; Gargiulo et al., 2009; Ohkawa et al., 2012). On the contrary, restriction endonucleases have been extensively used to digest the chromatin, presumably in an unbiased manner, in proximity ligation based techniques like 3C, 4C, 5C and HiC (Dekker et al., 2002; Dostie et al., 2006; Lieberman-Aiden et al., 2009; Zhao et al., 2006). In techniques like 3C, which has limited scope in terms of regions to be tested for spatial interactions, the efficiency of restriction digestion of the regions of interest can be tested and controlled. For high throughput assays like HiC, data on genome-wide assessment of digestion efficiency is rarely seen in the research articles and the associated supplementary materials. We showed that despite prolonged incubation with restriction endonuclease, the condensed nature of heterochromatin significantly limits its visibility to restriction endonucleases. As a result, interactions within heterochromatin and the ones impinging onto heterochromatin are under-represented in the current HiC datasets. This bias was not corrected by the methods that define the potential sources of bias explicitly. Widely used iterative correction method scales the contact matrices for equal visibility of each locus, without defining the source of bias at first place(Imakaev et al., 2012). We observed that the iterative correction method balanced the HiC signals in condensed and decondensed regions of the contact matrix locally. However, there were two issues with that: 1) It raised the long-range signals impinging from the condensed domains inadvertently. To mitigate this effect, we suggested a distance-dependent corrective step to be added post ICE-correction; 2) It only raised the obscure signals in the HiC matrix uniformly and did not resolve the structured sub-TAD pattern of the condensed domains as observed through GAM. This suggested that computational methods have their own limitations in resolving the obscure signals and HiC might need appropriate experimental refinement to resolve the condensed chromatin domains at sub-TAD levels.

Williamson et al had earlier reported discrepancy between 5C/HiC and the DNA FISH results concerning the condensation and decondensation of Hoxd locus during embryonic stem cell differentiation. While 5C/HiC largely showed that the locus remained condensed, DNA FISH clearly suggested that the locus decondensed upon differentiation (Williamson et al., 2014). We suggest that the biased efficiency of restriction digestion of condensed and decondensed forms of the locus might underlie such interpretations.

While we have shown that the biased visibility in HiC datasets can be largely addressed using ICE with an additional corrective step, the uncorrected bias itself is advantageous to explore another layer of chromatin organization. Differential visibility of chromatin domains help inferring dynamics of condensed and decondensed states of chromatin. Towards this, our analysis on developmental regulation of condensed and decondensed chromatin domains serve as a proof of principle. Coordinated changes in restriction enzyme accessibility and the epigenetic states of the genes contended against the arguments dismissing the observed bias merely as bias of trivial nature. The active chromatin marks exhibited shift towards the axis that represented the decondensed state of the involved domain in mESC or NPC, however the repressive H3K9me3 mark showed relatively subtle shift only. This was coherent with the earlier reports that suggested only subtle changes in H3K9 tri-methylation profiles during mESC differentiation (Filion and van Steensel, 2010; Lienert et al., 2011). We instead observed that the enrichment of polycomb associated proteins Suz12 and Ezh2 exhibited shift towards the axis that represented condensed state of chromatin domains (Figure S5h), suggesting that the non-LAD condensed domains uncovered in this study were likely representative of polycomb repressed chromatin. Polycomb association was also supported by the significant overlap of genes exhibiting decondesation during ESC-to-NPC transition with the Suz12 targets, Eed targets, PRC12 targets, and the targets of bivalent histone modifications (Figure S6b, right panel). Dynamical changes in the restriction enzyme accessibility were also observed during synthetic manipulation of the genome organization. Lamin deletion in the genome is associated with decondensation of LADs (Zheng et al, biorxiv 2018). We readily captured the decondensation of LADs in Lamin knock out cells from the HiC reads itself.

To identify the sequence features of domain boundaries, we tested the enrichment of several different genomic and epigenomic attributes (Figure 4b, S5c). We observed the enrichment of CTCF, RAD21, TOP2b, YY1 binding sites and repeat elements like MIR at boundaries, which are well known boundary elements of chromatin domains (Cuddapah et al., 2009; Heidari et al., 2014; Uuskula-Reimand et al., 2016; Wang et al., 2015). These observations collectively reinforce our claim on biological authenticity of condensed and decondensed domains identified outside the LAD regions.

A major criticism of our observations could be the proposal that potential source of the observed bias was differential restriction digestion. It can be argued that restriction digestion could be uniform, but the differential ligation could have caused the biased visibility. We do not entirely rule out the possibility that the restriction endonuclease nicked the DNA in condensed domains, but ligase enzyme failed to ligate the digested DNA due to stiffness and steric hindrance caused by nucleosomes and heterochromatin proteins in the condensed domains. However, we counter-argue that this may not be the case because when we plotted the HiC read counts against those of restriction digested chromatin, we observed a linear scaling, while there should have been bias towards the axis of restriction digested chromatin because of presumed efficient digestion and poor ligation in HiC experiments. We also proposed that the inefficient ligation of digested condensed chromatin can largely be a property of *in-situ* HiC and not the *in-solution* HiC, which involves dilution of digested chromatin and condensed chromatin is likely to exhibit relatively loose conformation owing to the usage of detergent and the heat during dilution step. We, therefore, compared the read counts of *in-situ* HiC and *in-solution* HiC datasets obtained from the same study. As shown in the figure S3c, we found good correlation between the two, while bias should have been reflected towards the axis representing reads from *in-solution* HiC assuming efficient digestion but poor ligation *in-situ* HiC. We thus proposed that the biased visibility in HiC datasets is caused by biased restriction digestion and not the ligation.

## Conclusion

Taken together, we highlighted a significant bias in the visibility of condensed and decondensed chromatin domains in HiC datasets attributing to non-uniform digestion of chromatin through restriction endonuclease. The existing computational methods failed to correct this bias appropriately and needed additional corrective measures. Finally, we showed that the repurposing of digestion bias was instrumental in deciphering another layer of gene-regulation through the dynamics of chromatin condensation and decondensation.

## Methods

We did not use any statistical method to predetermine sample size. We did not randomized any experiment. We were not blinded to allocations during experiments and outcome assessment. We mentioned the sample sizes and statistical tests wherever applicable. Source of each dataset is listed in Table S1

### Processing of HiC datasets and correction of 1D read counts

Wherever processed HiC data was not available, we obtained the SRA files and converted into fastq files using NCBI SRA toolkit. We implemented HiCUP package (http://www.bioinformatics.babraham.ac.uk/projects/hicup/) to process the HiC reads. We also used an in-house pipeline to process the HiC reads through bowtie2 (http://bowtie-bio.sourceforge.net/bowtie2/index.shtml). In brief, we mapped the paired-end reads on indexed mm10 assembly of mouse genome. Reads were then paired and filtered for invalid and duplicated reads. To make one-dimensional chromosomal tracks of HiC data, we mixed together the HiCUP processed paired-end reads and calculated the read counts for 10kb bins of genome. To normalize the known systematic biases in HiC contact maps, we used HiC-Norm (http://www.people.fas.harvard.edu/~junliu/HiCNorm/) and ICE packages (https://github.com/mirnylab). Table S1 marks the details of HiC datasets and their processing details. We obtained the bedgraph files (mm9) for *in-situ* digested chromatin and *in-solution* digested naked DNA from GSE51821. The files were converted into mm10 assembly using UCSC’s liftover utility (http://genome.ucsc.edu/cgi-bin/hgLiftOver). Sequence reads were binned into 10kb bins genome-wide. Relative density of restriction enzyme sites might influence the 1D read counts obtained after HiCUP processing. We, therefore, removed the RE density associated bias in the read counts. We used the residuals of read counts after loess regression against RE density of genomic bins. 1D read counts are also biased due to varying GC content of genomic regions. GC rich domains are readily captured in sequencing reactions as compared to GC-poor regions. We removed this bias by calculating residuals of RE-corrected read counts through loess regression against GC content of corresponding genomic bins. The final corrected read count had no scaling against RE density and GC content of the genomic bins as shown in the Figure S1a-b.

### Domain calling

We scaled the 1D HiC read counts using Z-score and identified the condensed and decondensed domains using the strategy given by Guelen et al (Guelen et al., 2008). In short, read counts were first binarized as +1 and −1 depending upon whether the values were positive of negative in the Z-scores. The domain boundaries were identified by subtracting the average of 20 windows on either side of uniformly distributed (per 10 kb) reference points. We determined a cut-off on this value through randomization of the read counts in the genome and keeping the false discovery rate to <5%. By calculating the relative proportion of positive and negative values in each inter-boundary regions, we demarcated condensed and decondensed domains. We set the minimal proportion of either positive or negative values to 0.8 in order to classify the domains as decondensed and condensed respectively.

### Analysis of constitutive LADs and constitutive inter-LADs

We downloaded 3843 LAD regions of mouse ES cells, neuronal progenitor cells and astrocytes from GSE17051. We classified LADs as constitutive LADs (cLADs) and constitutive inter-LADs (ciLADs) by comparing the genome coordinates of LADs in mESC, NPC and astrocytes. Similarly, 605 cLADs in human were obtained by comparing LAD coordinates in IMR90 and heterochromatin domain coordinates in h1ESC and K562 cell-lines from ENCODE (http://www.encodeproject.org/comparative/chromatin/). To analyse the alteration in the conformation of cLAD domains, we obtained the Lamin KO HiC data for mESCs, binned the reads into 10 kb bins and used log ratio of Lamin KO to WT for our analyses.

### Analysis of polytene and normal diploid HiC in Drosophila

We downloaded HiC SRA files from GSE72510 (polytene, dm6) and GSE63518 (normal diploid Kc167, dm6) and processed using HiCUP pipeline. We binned the reads at 5kb resolution for RE-density and GC correction of the read counts. We downloaded polytene TADs from Eagen et al (2015) and lifted over the coordinates to dm6 assembly. We mapped 5kb bins to polytene TADs and considered those inside TADs as polytene band bins and rest as inter-band. We generated raw contact maps at a 5kb resolution and normalized using HiC-norm and ICE using python *mirnylab* package.

### Allele-specific HiC analysis of X-chromosome

We downloaded allele-specific valid HiC pairs of brain and patski cells from GSE68992. We removed the reads mapping to both references and binned the allele-specific reads at 20kb resolution to obtain one dimensional read counts. We corrected the read counts for GC content using loess regression as earlier. To visualize the interactions, we generated raw and ICE corrected contact maps at 100kb. HiCNorm does not suit to DNase-HiC data due to the usage of DNase instead of restriction endonuclease, and hence not used.

### Analysis of histone modification and CTCF ChIP-seq datasets

Source of ChIP-Seq datasets are given in Table S1. Then, we binned the reads at 10kb resolution, and quantile normalized using the R package preprocessCore (https://github.com/bmbolstad/preprocessCore). We generated median aggregate plots by aligning all boundaries that have at least 200kb long open/close on their either sides. We used R-package *ggplot2* (https://ggplot2.tidyverse.org/) to scatterplot the data.

### Distance Sorted Contact Optimization (DiSCO)

We optimized the raw, HiCNorm-corrected and ICE-corrected HiC data for the distance dependent bias in the interaction frequencies of condensed and decondensed domains. We balanced the interaction frequencies in condensed and decondensed domains (as identified from corrected 1D read counts) using quantile normalization of the distance sorted mean frequency of interactions for each 10kb genomic bin-pair. We then adjusted the values in the contact matrices using the quantile normalized mean, the Z-score and the standard deviations of each distance bin-pair.

### Data availability

The bedgraph files for the corrected read counts of all the HiC datasets are available at following link: https://bitbucket.org/ken_at_keerthivasan/compaction_from_hic/downloads/

### Funding

This work was supported by Department of Biotechnology (DBT-India),

(BT/PR13596/GET/119/31/2015; BT/PR8688/AGR/36/755/2013;

BT/PR16366/BID/7/598/2016), Indian Council of Medical

Research (F.No.90/09/2012/SCRT(TF)/BMS) and SERB (EMR/2015/001681).

## Acknowledgements

KRC and PKG acknowledge ICMR and CSIR respectively for their fellowship support.

