## Supplementary Tables and Figures for "Repurposing the biased visibility in HiC datasets to mark dynamically regulated condensed and decondensed chromatin states genome-wide"

#### Supplementary Table and Figure legends

**Table S1.** Details of the datasets used in the study.

**Figure S1.** Related to figure 1. (a) Loess correction for the negative scaling of raw read counts against the restriction site (RE) density in 10 Kb genomic bins. Left panel represents data before loess correction and right panel after loess correction of read counts against RE-density (b) Loess correction for the positive scaling of RE-corrected read counts against the GC content of 10 Kb genomic bins. First panel shows scatter plot of GC content vs. RE-corrected read counts. Second panel shows scatter plot of GC content vs. GC- and RE-corrected read counts. Third panel represents RE-density vs. GC- and RE-corrected read counts. Fourth panel shows scaling of GC- and RE-corrected read counts against the raw read counts. (c) Size distribution of domains identified through analysis of corrected read counts. Plotted are the mean values with the standard error bars (d) Genomic coverage of condensed domains within constitutive LAD and constitutive inter-LAD regions. Shown are the pie charts of 10Kb bins mapping to cLAD and ciLADs in different datasets.

**Figure S2.** Related to Figure 1. (a) Distributions of raw and corrected read count in cLAD and ciLADs across different *in-situ* HiC datasets in mouse. (b) Distributions of raw and corrected read count in cLAD and ciLADs across different *in-situ* HiC datasets in human. We calculated p-values using two-tailed Mann Whitney U test.

**Figure S3.** Related to Figure 1. (a) Distributions of bowtie-processed raw and corrected read counts in cLAD and ciLADs across *in-situ* HiC datasets of mESC, NPC and CN cells. We calculated p-values using two-tailed Mann Whitney U test. (b) Side-by-side comparison of raw and corrected read counts mapping to cLAD and ciLADs in *in-situ* and *in-solution* (dilution) HiC datasets obtained for the same cells (mouse fetal liver) from the same study. (c) Scatter plot of corrected read counts obtained from *in-situ* and *in-solution* HiC datasets.

**Figure S4**. Related to Figure 2. (a) Distribution of interaction frequencies of decondensed-to-decondensed and condensed-to-condensed interactions as a function of genomic distance in the raw, HiCNorm-corrected and ICE-corrected HiC, and GAM datasets. Upper and lower panels show plots without and with DiSCO corrections respectively. Both axes are log10 tranformed and y-axis was further scaled from 0 to 1 for comparison across plots. (b) Distribution of ICE+DiSCO corrected 1D read counts in the condensed and decondensed domains. (c) Additional examples comparing the corrected 1D read counts and the contact matrices of raw, HiCNorm-corrected, ICE-corrected HiC, and the GAM datasets. (d-e) Additional examples comparing the contact matrices of raw, HiCNorm-corrected, and ICE-corrected HiC datasets with and without DiSCO correction.

**Figure S5.** Related to Figure4. (a) Scatter plots of corrected read counts in mESC vs. NPC and in NPC vs. CN. (b) Enrichment of histone modifications around boundary between decondensed and condensed domains in mouse cortical neurons (CN). (c) Enrichment of various genomic attributes around domain boundaries and domain centers. (d) Distribution of condensed and decondensed states of chromatin domains during mESC to NPC differentiation. (e-g) Examples of histone modification profiles around ciLAD condensed and decondensed domains in mESC and NPC. (h) Scatter plots for enrichment of Polycomb proteins in mESC vs. NPC with respect to constitutive and cell-type specific status of the ciLAD condensed and decondensed chromatin domains.

**Figure S6.** Related to Figure 4. (a) Enrichment of Gene Ontology Process terms among the genes exhibiting condensation (left) and decondensation (right) during ESC-to-NPC transition. Shown are the top 30 terms through ToppGene Suite. Nervous system associated terms are highlighted in brown colour. (b) Significance of overlap between MSigDB gene sets and the genes exhibiting condensation (left) and decondensation (right) during ESC-to-NPC transition. Shown are the top 30 terms through Gene Set Enrichment Analysis (GSEA). Polycomb associated terms are highlighted in brown colour. Vertical dashed line in each plot marks FDR of 0.01.

**Table S1**

| Accession | Cell-type | Experiment | RE | Processing |
|---|---|---|---|---|
| GSE96107 | ES, NP, CN | In-situ HiC | DpnII | HiCUP/bowtie |
| GSE59027 | ES, NP, CN | In-solution HiC | NcoI, | HiCUP/bowtie |
| | | In-solution HiC | HindIII | |
| GSE72510 | Polytene | Tethered HiC | DpnII | HiCUP/bowtie |
| GSE63518 | Diploid Kc167 | In-solution HiC | DpnII | HiCUP/bowtie |
| GSE89520 | ES (lamin KO) | In-situ HiC | BglII | Pre-processed |
| GSE68992 | Brain, Patski | DNase-HiC | DNase | Pre-processed |
| GSE72697 | ES, NP | In-solution HiC | HindIII | Pre-processed |
| GSE70181 | Liver | In-situ/in-solution HiC | HindIII | Pre-processed |
| GSE74055 | ES | In-situ HiC | MboI | Pre-processed |
| GSE86150 | ES | In-situ HiC | MboI | Pre-processed |
| GSE99991 | NP | In-situ HiC | HindIII | Pre-processed |
| GSE94452 | ES | In-situ HiC | MboI | Pre-processed |
| GSE107282 | Patski | DNase-HiC | DNase | Pre-processed |
| GSE63525 | GM12878, IMR90, HMEC, | In-situ/in-solution HiC | HindIII, DpnII (GM12878) | Pre-processed |
| | HUVEC | | | |
| GSE80820 | GM12878 | In-situ HiC | MboI | Pre-processed |
| GSE107148 | HSPC | In-situ HiC | MboI | Pre-processed |
| GSE98671 | ES | In-solution HiC | HindIII | Pre-processed |
| GSE104129 | Liver | In-situ HiC | MboI | HiCUP |
| GSE103477 | MDM | In-situ HiC | MboI | Pre-processed |
| GSE80280 | ES | Single cell HiC | MboI | HiCUP |
| ENCSR032JUI | ES | H3K4me1 ChIP-seq | | Pre-processed |
| ENCSR000CGQ | | H3K27ac ChIP-seq | | Pre-processed |
| ENCSR000CGO | | H3K4me3 ChIP-seq | | Pre-processed |
| ENCSR253QPK | | H3K36me3 ChIP-seq | | Pre-processed |
| ENCSR857MYS | | H3K9me3 ChIP-seq | | Pre-processed |
| ENCSR059MBO | | H3K27me3 ChIP-seq | | Pre-processed |
| GSE96107 | | CTCF ChIP-seq | | Pre-processed |
| GSE96107 | NP | H3K4me1 ChIP-seq | | Pre-processed |
| | | H3K27ac ChIP-seq | | Pre-processed |
| | | H3K4me3 ChIP-seq | | Pre-processed |
| | | H3K36me3 ChIP-seq | | Pre-processed |
| | | H3K9me3 ChIP-seq | | Pre-processed |
| | | H3K27me3 ChIP-seq | | Pre-processed |
| | | CTCF ChIP-seq | | Pre-processed |
| GSE96107 | CN | H3K4me1 ChIP-seq | | Pre-processed |
| | | H3K27ac ChIP-seq | | Pre-processed |
| | | H3K4me3 ChIP-seq | | Pre-processed |
| | | H3K36me3 ChIP-seq | | Pre-processed |
| | | H3K9me3 ChIP-seq | | Pre-processed |
| | | H3K27me3 ChIP-seq | | Pre-processed |
| | | CTCF ChIP-seq | | Pre-processed |
| GSE31786 | ES | YY1 ChIP-seq | | Pre-processed |
| GSE90994 | ES | Rad21 ChIP-seq | | Pre-processed |
| E-MTAB-3587 | liver | Top2b ChiP-seq | | Pre-processed |
| Repeats | Mouse | UCSC browser | | Pre-processed |
| GSE96107 | ES | RNA-seq | | Pre-processed |
| | NP | | | Pre-processed |
| | CN | | | Pre-processed |

### Figure S1

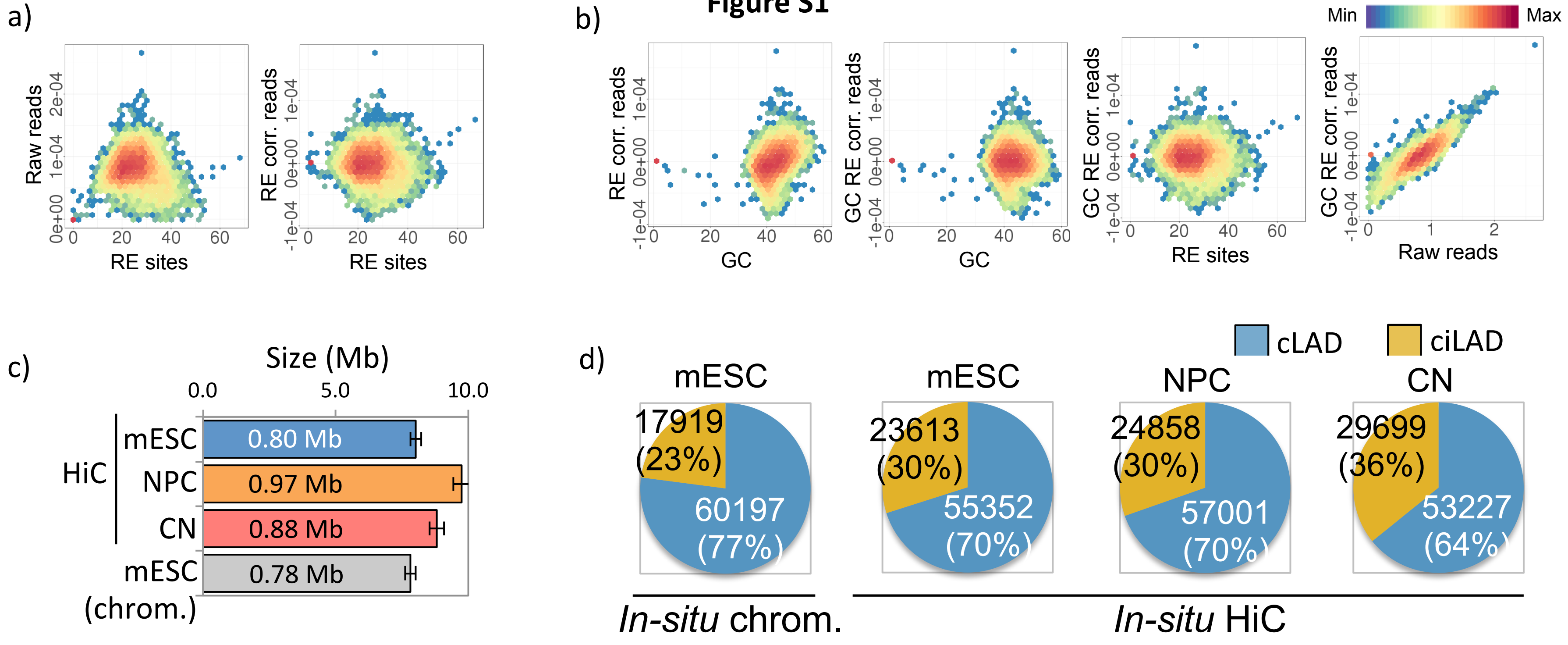

a)
Raw reads
RE corr. reads
RE sites
b)
Min
Max
RE corr. reads
GC RE corr. reads
GC
RE sites
Raw reads
c)
Size (Mb)
0.0
5.0
10.0
HiC
mESC
0.80 Mb
NPC
0.97 Mb
CN
0.88 Mb
mESC (chrom.)
0.78 Mb
d)
cLAD
ciLAD
mESC
17919 (23%)
60197 (77%)
mESC
23613 (30%)
55352 (70%)
NPC
24858 (30%)
57001 (70%)
CN
29699 (36%)
53227 (64%)
In-situ chrom.
In-situ HiC

### Figure S2

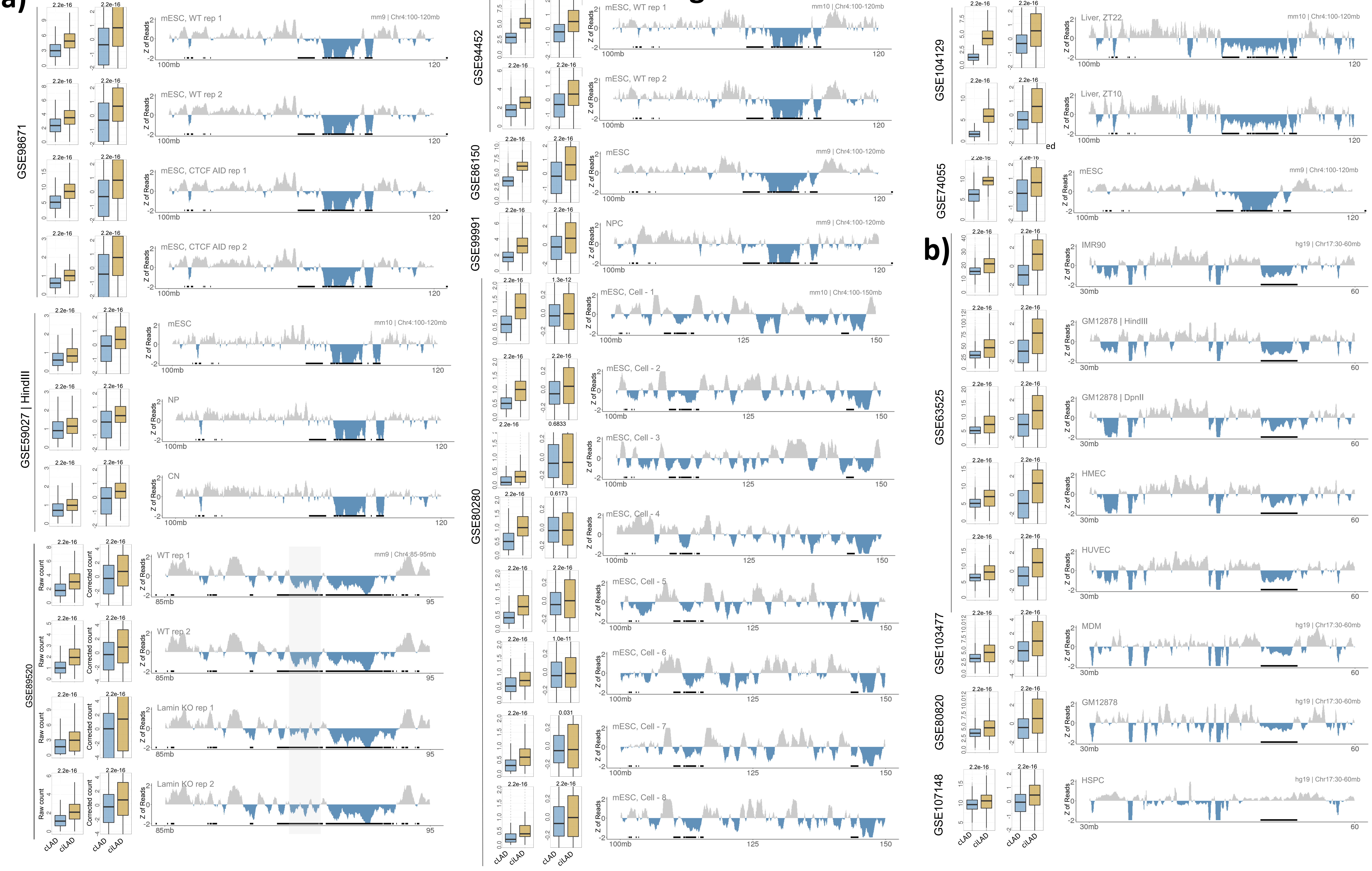

a)
b)
Raw
Corrected
GSE98671
GSE59027 | HindIII
GSE89520
GSE94452
GSE86150
GSE99991
GSE80280
GSE104129
GSE74055
GSE63525
GSE103477
GSE80820
GSE107148
mESC, WT rep 1
mESC, WT rep 2
mESC, CTCF AID rep 1
mESC, CTCF AID rep 2
mESC
NP
CN
WT rep 1
WT rep 2
Lamin KO rep 1
Lamin KO rep 2
NPC
mESC, Cell - 1
mESC, Cell - 2
mESC, Cell - 3
mESC, Cell - 4
mESC, Cell - 5
mESC, Cell - 6
mESC, Cell - 7
mESC, Cell - 8
Liver, ZT22
Liver, ZT10
IMR90
GM12878 | HindIII
GM12878 | DpnII
HMEC
HUVEC
MDM
GM12878
HSPC
mm9 | Chr4:100-120mb
mm10 | Chr4:100-120mb
mm9 | Chr4:85-95mb
mm10 | Chr4:100-150mb
hg19 | Chr17:30-60mb
Z of Reads
Raw count
Corrected count
cLAD
ciLAD
100mb
120
85mb
95
125
150
30mb
60
2.2e-16
1.3e-12
0.6833
0.6173
1.0e-11
0.031

### Figure S3

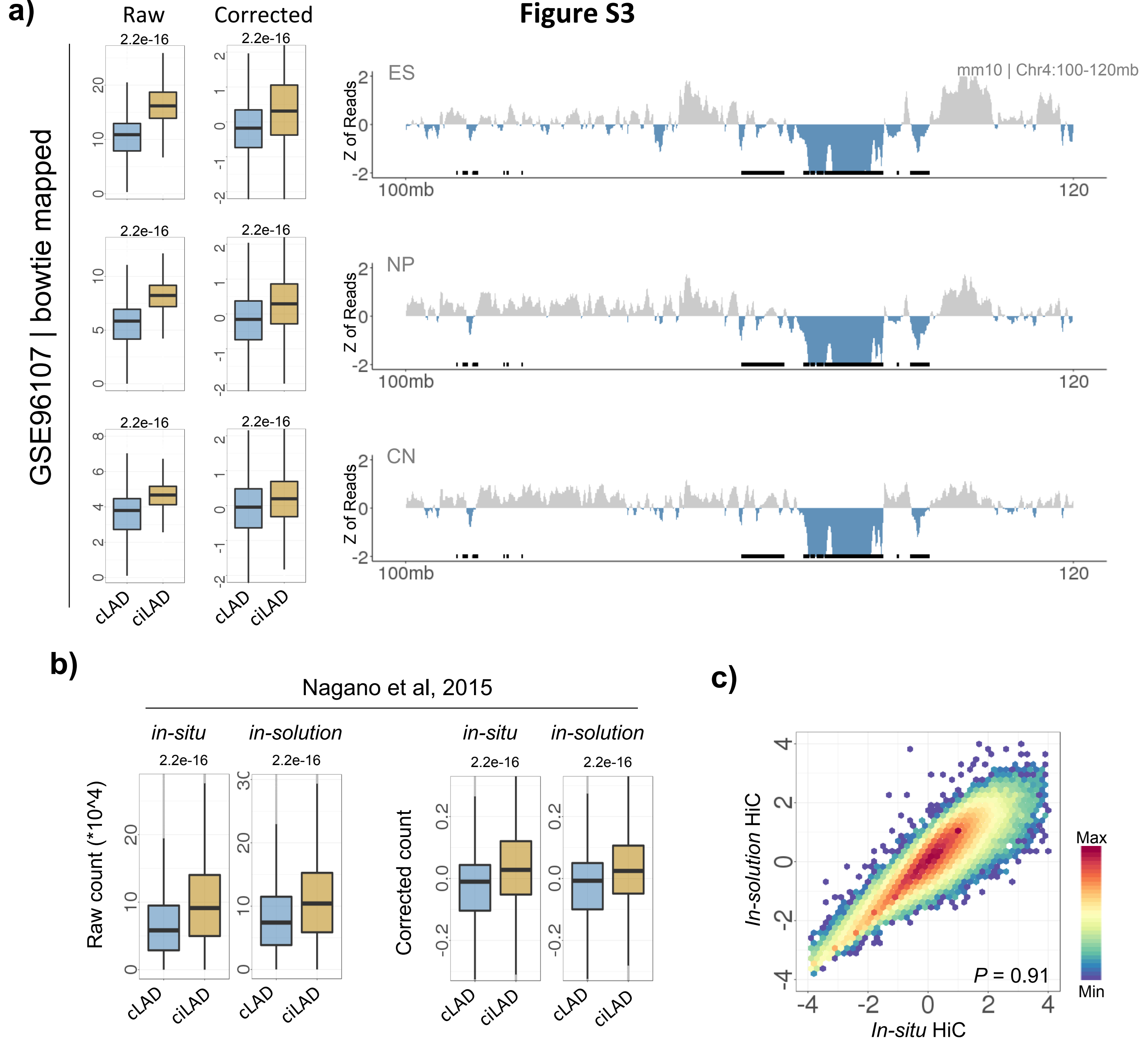
a)
Raw
Corrected
GSE96107 | bowtie mapped
2.2e-16
cLAD
ciLAD
ES
NP
CN
mm10 | Chr4:100-120mb
Z of Reads
100mb
120
b)
Nagano et al, 2015
in-situ
in-solution
Raw count (*10^4)
Corrected count
c)
In-solution HiC
In-situ HiC
P = 0.91
Max
Min

### Figure S4

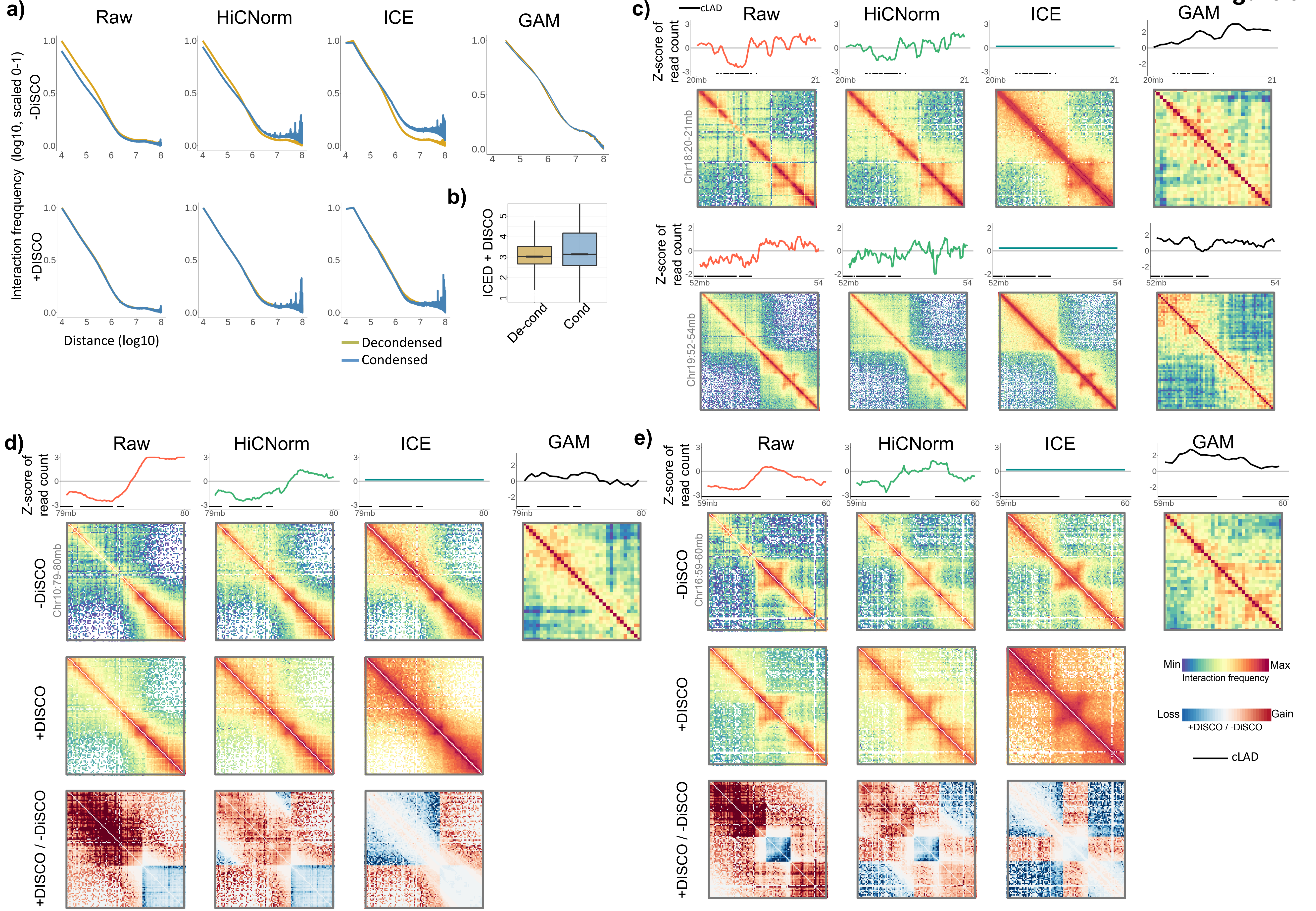
a)
Raw
HiCNorm
ICE
GAM
Interaction freqquency (log10, scaled 0-1)
-DiSCO
+DISCO
Distance (log10)
Decondensed
Condensed
b)
ICED + DISCO
De-cond
Cond
c)
cLAD
Z-score of read count
20mb
21
Chr18:20-21mb
52mb
54
Chr19:52-54mb
d)
79mb
80
-DiSCO
Chr10:79-80mb
+DISCO
+DISCO / -DiSCO
e)
59mb
60
Chr16:59-60mb
Min
Max
Interaction frequency
Loss
Gain
+DISCO / -DiSCO
cLAD

### Figure S5

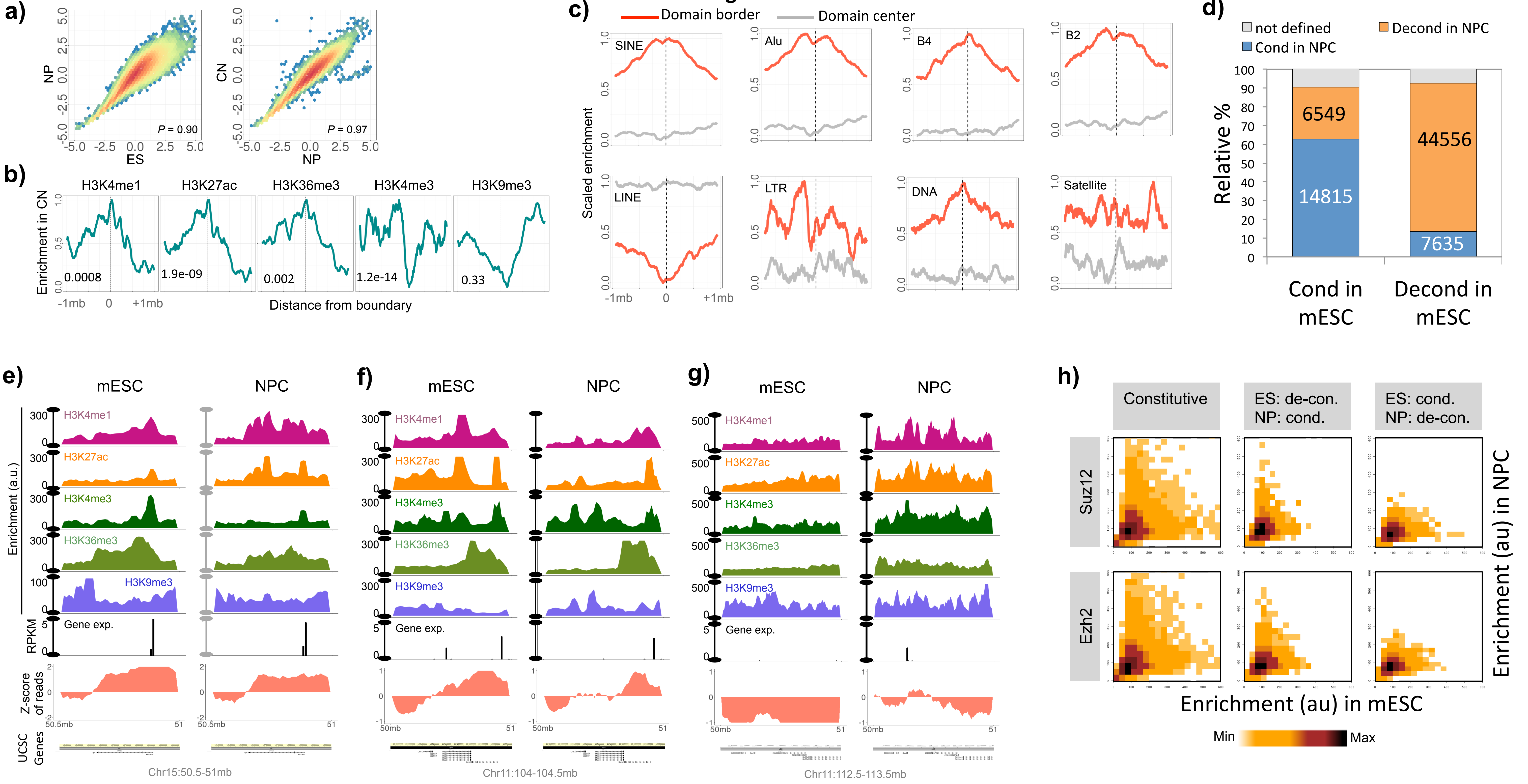

a)
NP
ES
P = 0.90
CN
NP
P = 0.97
b)
H3K4me1
H3K27ac
H3K36me3
H3K4me3
H3K9me3
Enrichment in CN
0.0008
1.9e-09
0.002
1.2e-14
0.33
-1mb
0
+1mb
Distance from boundary
c)
Domain border
Domain center
Scaled enrichment
SINE
Alu
B4
B2
LINE
LTR
DNA
Satellite
-1mb
0
+1mb
d)
not defined
Cond in NPC
Decond in NPC
Relative %
6549
14815
44556
7635
Cond in mESC
Decond in mESC
e)
mESC
NPC
Enrichment (a.u.)
H3K4me1
H3K27ac
H3K4me3
H3K36me3
H3K9me3
Gene exp.
RPKM
Z-score of reads
UCSC Genes
50.5mb
51
Chr15:50.5-51mb
f)
mESC
NPC
H3K4me1
H3K27ac
H3K4me3
H3K36me3
H3K9me3
Gene exp.
50mb
51
Chr11:104-104.5mb
g)
mESC
NPC
H3K4me1
H3K27ac
H3K4me3
H3K36me3
H3K9me3
Gene exp.
50mb
51
Chr11:112.5-113.5mb
h)
Constitutive
ES: de-con.
NP: cond.
ES: cond.
NP: de-con.
Suz12
Ezh2
Enrichment (au) in NPC
Enrichment (au) in mESC
Min
Max

Figure S6

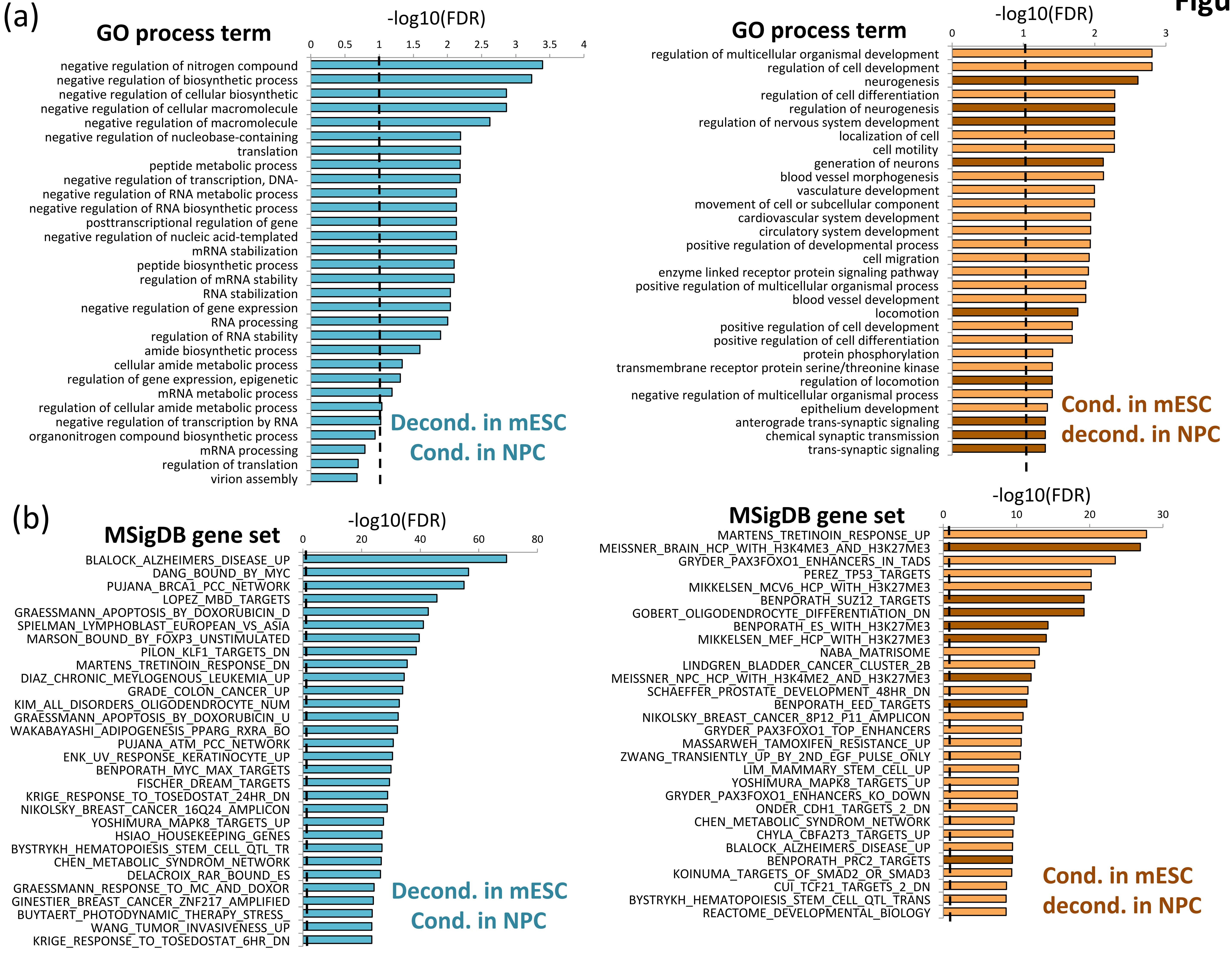
(a)
GO process term
-log10(FDR)
0 0.5 1 1.5 2 2.5 3 3.5 4
negative regulation of nitrogen compound
negative regulation of biosynthetic process
negative regulation of cellular biosynthetic
negative regulation of cellular macromolecule
negative regulation of macromolecule
negative regulation of nucleobase-containing
translation
peptide metabolic process
negative regulation of transcription, DNA-
negative regulation of RNA metabolic process
negative regulation of RNA biosynthetic process
posttranscriptional regulation of gene
negative regulation of nucleic acid-templated
mRNA stabilization
peptide biosynthetic process
regulation of mRNA stability
RNA stabilization
negative regulation of gene expression
RNA processing
regulation of RNA stability
amide biosynthetic process
cellular amide metabolic process
regulation of gene expression, epigenetic
mRNA metabolic process
regulation of cellular amide metabolic process
negative regulation of transcription by RNA
organonitrogen compound biosynthetic process
mRNA processing
regulation of translation
virion assembly
Decond. in mESC
Cond. in NPC
GO process term
-log10(FDR)
0 1 2 3
regulation of multicellular organismal development
regulation of cell development
neurogenesis
regulation of cell differentiation
regulation of neurogenesis
regulation of nervous system development
localization of cell
cell motility
generation of neurons
blood vessel morphogenesis
vasculature development
movement of cell or subcellular component
cardiovascular system development
circulatory system development
positive regulation of developmental process
cell migration
enzyme linked receptor protein signaling pathway
positive regulation of multicellular organismal process
blood vessel development
locomotion
positive regulation of cell development
positive regulation of cell differentiation
protein phosphorylation
transmembrane receptor protein serine/threonine kinase
regulation of locomotion
negative regulation of multicellular organismal process
epithelium development
anterograde trans-synaptic signaling
chemical synaptic transmission
trans-synaptic signaling
Cond. in mESC
decond. in NPC
(b)
MSigDB gene set
-log10(FDR)
0 20 40 60 80
BLALOCK_ALZHEIMERS_DISEASE_UP
DANG_BOUND_BY_MYC
PUJANA_BRCA1_PCC_NETWORK
LOPEZ_MBD_TARGETS
GRAESSMANN_APOPTOSIS_BY_DOXORUBICIN_D
SPIELMAN_LYMPHOBLAST_EUROPEAN_VS_ASIA
MARSON_BOUND_BY_FOXP3_UNSTIMULATED
PILON_KLF1_TARGETS_DN
MARTENS_TRETINOIN_RESPONSE_DN
DIAZ_CHRONIC_MEYLOGENOUS_LEUKEMIA_UP
GRADE_COLON_CANCER_UP
KIM_ALL_DISORDERS_OLIGODENDROCYTE_NUM
GRAESSMANN_APOPTOSIS_BY_DOXORUBICIN_U
WAKABAYASHI_ADIPOGENESIS_PPARG_RXRA_BO
PUJANA_ATM_PCC_NETWORK
ENK_UV_RESPONSE_KERATINOCYTE_UP
BENPORATH_MYC_MAX_TARGETS
FISCHER_DREAM_TARGETS
KRIGE_RESPONSE_TO_TOSEDOSTAT_24HR_DN
NIKOLSKY_BREAST_CANCER_16Q24_AMPLICON
YOSHIMURA_MAPK8_TARGETS_UP
HSIAO_HOUSEKEEPING_GENES
BYSTRYKH_HEMATOPOIESIS_STEM_CELL_QTL_TR
CHEN_METABOLIC_SYNDROM_NETWORK
DELACROIX_RAR_BOUND_ES
GRAESSMANN_RESPONSE_TO_MC_AND_DOXOR
GINESTIER_BREAST_CANCER_ZNF217_AMPLIFIED
BUYTAERT_PHOTODYNAMIC_THERAPY_STRESS_
WANG_TUMOR_INVASIVENESS_UP
KRIGE_RESPONSE_TO_TOSEDOSTAT_6HR_DN
Decond. in mESC
Cond. in NPC
MSigDB gene set
-log10(FDR)
0 10 20 30
MARTENS_TRETINOIN_RESPONSE_UP
MEISSNER_BRAIN_HCP_WITH_H3K4ME3_AND_H3K27ME3
GRYDER_PAX3FOXO1_ENHANCERS_IN_TADS
PEREZ_TP53_TARGETS
MIKKELSEN_MCV6_HCP_WITH_H3K27ME3
BENPORATH_SUZ12_TARGETS
GOBERT_OLIGODENDROCYTE_DIFFERENTIATION_DN
BENPORATH_ES_WITH_H3K27ME3
MIKKELSEN_MEF_HCP_WITH_H3K27ME3
NABA_MATRISOME
LINDGREN_BLADDER_CANCER_CLUSTER_2B
MEISSNER_NPC_HCP_WITH_H3K4ME2_AND_H3K27ME3
SCHAEFFER_PROSTATE_DEVELOPMENT_48HR_DN
BENPORATH_EED_TARGETS
NIKOLSKY_BREAST_CANCER_8P12_P11_AMPLICON
GRYDER_PAX3FOXO1_TOP_ENHANCERS
MASSARWEH_TAMOXIFEN_RESISTANCE_UP
ZWANG_TRANSIENTLY_UP_BY_2ND_EGF_PULSE_ONLY
LIM_MAMMARY_STEM_CELL_UP
YOSHIMURA_MAPK8_TARGETS_UP
GRYDER_PAX3FOXO1_ENHANCERS_KO_DOWN
ONDER_CDH1_TARGETS_2_DN
CHEN_METABOLIC_SYNDROM_NETWORK
CHYLA_CBFA2T3_TARGETS_UP
BLALOCK_ALZHEIMERS_DISEASE_UP
BENPORATH_PRC2_TARGETS
KOINUMA_TARGETS_OF_SMAD2_OR_SMAD3
CUI_TCF21_TARGETS_2_DN
BYSTRYKH_HEMATOPOIESIS_STEM_CELL_QTL_TRANS
REACTOME_DEVELOPMENTAL_BIOLOGY
Cond. in mESC
decond. in NPC
